# A compromised gsdf signaling leads to gamatogenesis confusion and subfertility in medaka

**DOI:** 10.1101/238436

**Authors:** Guijun Guan, Shumei Xu, Anning Guo, Xiaomiao Zhao, Yingqing Zhang, Kaiqing Sun, Yi Kang, Yuyang Chang, Xiaowen Wu, Liangbiao Chen

**Author notes:** These authors share the first authorship (Guijun Guan, Shumei Xu). To whom correspondence should be addressed: **Dr. Guijun Guan**, College of Fisheries and Life Sciences, Shanghai Ocean University, HuchengHuan Road 999, Shanghai 201306, China; **Dr. Liangbiao Chen**, College of Fisheries and Life Sciences, Shanghai Ocean University, HuchengHuan Road 999, Shanghai 201306, China.

## Abstract

**Summary statement:** Gsdf signals trigger the gamatogenesis, alter the somatic expression of Fsh/Lh receptors and brain type aromatase in medaka brain and gonad.

**Abstract:** Gonadal soma-derived factor (*gsdf*) and anti-Mullerian hormone (*amh*) are somatic male determinants in several species of teleosts, although the mechanisms by which they trigger the indifferent germ cells into the male pathway remain unknown. This study aimed to decipher the roles of *gsdf/amh* in directing the sexual fate of germ cells using medaka as a model. Transgenic lines (Tg^cryG^) that restrictively and persistently express a Gsdf-Gfp fusion protein in the lens and the hypothalamus-pituitary-gonad (HPG) axis, were generated under the control of a mouse γF-crystallin promoter. A high frequency (44.4%) of XX male sex reversals was obtained in Tg^cryG^ lines, indicating that signals of gsdf-expressing cells in HPG were enough for the spermatogenesis activation in the genetic females. Furthermore, all Tg^cryG^ XY individuals with endogenous *gsdf* depletion (named *Sissy*) displayed intersex (100%) with enlarged ovotestis in contrast to a giant ovary developed in XY *gsdf* deficiency. The heterogeneous expression of *gsdf* led to the confusion of gamatogenesis and ovotestis development, similar to some *hotei (amhr2*) mutants, suggests that the signaling balance of *gsdf/amh* is essential for proper gamatogenesis, maintaining sex steroid production and gonadotropin secretion, which are evolutionarily conserved across *phyla*.

## Introduction

Sex determination in lower vertebrates, particularly in teleost fish is highly diverse. They are liable to sex exchange in response to environmental elements, such as temperature, sex hormones, or chemical contaminants, even their sex is largely determined genetically(Bachtrog et al., 2014). In medaka (*Oryzias latipes*), male sex is determined under the influence of *<u>d</u>oublesex-<u>m</u>ab3* domain gene on <u>Y</u> chromosome (*dmy* gene) (Matsuda et al., 2002), but can be turnover by targeted disruption of other genes such as gonadal soma-derived factor (*gsdf*) which is important for testis development(Imai et al., 2015; Zhang et al., 2016). Medaka sex can also be artificially altered by steroid hormones, or by manipulating the number of germ cells (Nagahama et al., 2004). Therefore, medaka serves as an excellent vertebrate model for the study of sexual differentiation and plasticity.

Gsdf depletion not only resulted in XY feminization, but also severely restrained oocyte development at previtellogenic stage(Guan, 2017). Similar phenomena have been observed after targeted disruption of follicle-stimulating hormone (*fsh*)(Takahashi et al., 2016), *fsh* receptor (*fshr*)(Murozumi et al., 2014) or an anti-Müllerian hormone receptor II (*amhr2*) *hotei* mutant leading to female sterility(Morinaga et al., 2007). Fsh, Amh, androgens, and estradiol (E2) are essential in folliculogenesis in most vertebrates, but displays a wide range of effective varieties in spermatogenesis(Huhtaniemi, 2015). Fsh is required for ovarian folliculogenesis in mammals and fish, because *Fsh* knockout (*FshKO*) mice displays a blockade of folliculogenesis at the preantral follicle stage and leads to female infertility(Kumar, 2009). In contrast, both male mice and medaka were fertile with reduced testis size(Kumar et al., 1997). Disruption of *fshr* leads to the arrest of ovary development and turns on a small portion (10%) of male sex reversal in zebrafish and medaka (Murozumi et al., 2014; Zhang et al., 2015), but causes failure of normal Leydig cell development in mice(Baker et al., 2003). The expression of *Fsh* β-subunit can be detected from embryonic stages in mammals and fish(Weltzien et al., 2014), showing a sexual dimorphism of Fsh-expressing cells more in female medaka before hatching(Horie et al., 2014). Another gonadotropin Luteinizing hormone (Lh) shares a common glycoprotein α-subunit (GPA) with Fsh, but has a hormone-specific β-subunit separately from *fsh* in fish(Hsu et al., 2002). The embryological expression of Lh was also detectable at a late gastrulation stage using sensitive real-time polymerase chain reaction (PCR) and *lh–gfp* transgenic fish(Hildahl et al., 2012). Impairment of oocyte development and severe previtellogenic oocyte arrest have been observed with the depletion of *gsdf*, probably due to the dramatic decrease in the expression of *fshr* (Kaiqing Sun1#, 2017). However, the molecular events including TGF-β factors, gonadotropins, and their receptors during gonadal differentiation are still not fully understood.

Amh signaling is the key player in gonad development conserved from fish to mammals, which effects in the hypothalamic-pituitary-gonadal axis (HPG) as a target of gonadotropic Fsh in teleost fish(Pfennig et al., 2015). Amh and Gsdf share structural similarity in TGF-β domain, and also display a close correlation to the testicular differentiation of gonochoristic medaka or sex transformation of rice field eels (*Monopterus albus*), the hermaphrodite fish(Pfennig et al., 2015). The phenotype of *amhr2* mutant *hotei* is similar to *gsdf KO* because both display a hypertrophic ovary in XX and XY homozygotes, despite a half XY feminization but the other half XY normal males in *hotei*, in contrast to a complete feminization in *gsdf* deficiency(Zhang et al., 2016). Evidences from germline chimeras generated by germ cells transplantation between *hotei* and wild-type XY gonads demonstrated AMH signaling ruling germ cell proliferation indirectly via unknown signals(Nakamura et al., 2012). Gsdf is considered as one of this unknown signals, therefore this study aimed to investigate the effect of *gsdf* signaling in HPG axis, with a transgenic medaka expressing Gsdf and Egfp under the control of mouse γF-crystalline promoter(Vopalensky et al., 2010). The exogenous *gsdf* induced XX maleness and partially resumed spermatogenesis in the absence of endogenous *gsdf* (*gsdf*KO) otherwise, an ovary was developed, exactly replicated ovotestis appearance derived from a small portion of *hotei* mutants. Thus, the study provides novel insights of molecular mechanisms underlying the sexual differentiation of germ cells and gonadal development.

## Results

### Establishment of *gsdf* transgenic lines

Previous data revealed that the over-expression of *gsdf2AEgfp* driven by β-actin promoter in gonads led to XX-testis development in medaka(Zhang et al., 2016). A transgenic line was generated with the help of mouse γF-crystalline promoter (Tg^cryG^), in which *gsdf–gfp* was expressed in lens, brain, and gonad (Fig 1). EGFP in the backbone of p817-EGFP (Addgene) reporter construct was replaced with a gsdf2AEGFP fragment in the frame(Vopalensky et al., 2010). The schematic representation of the expression construct is shown in Figure 1a. Both circular and linear DNAs (digested with I-SceI) were microinjected into 1256 fertilized eggs at a one-cell stage in 4 individual experiments. Subsequently, 516 hatched eggs were obtained, and 276 grew to be reproductive mature adults. The total survival rate and transgenic efficiency are summarized in Table S1. Of these, 17 fish displayed GFP signals in their lens from the early embryo stage (Figure 1b–b’), which were retained until adulthood (Fig. 1c and 1d). All GFP-positive F0 fish were fertile, and three founder fish (one male and two females) passed GFP signals to their progeny after outcrossing with wild-type males and females, indicating that the transgenic expression of gsdf2AEGFP did not disturb gonadal development. Phenotypic sex was completely matched with their genotypic sex, indicating that no sex reversal occurred overriding the control of *dmy* as the male-determining factor in F0 generation.

**Figure 1.**
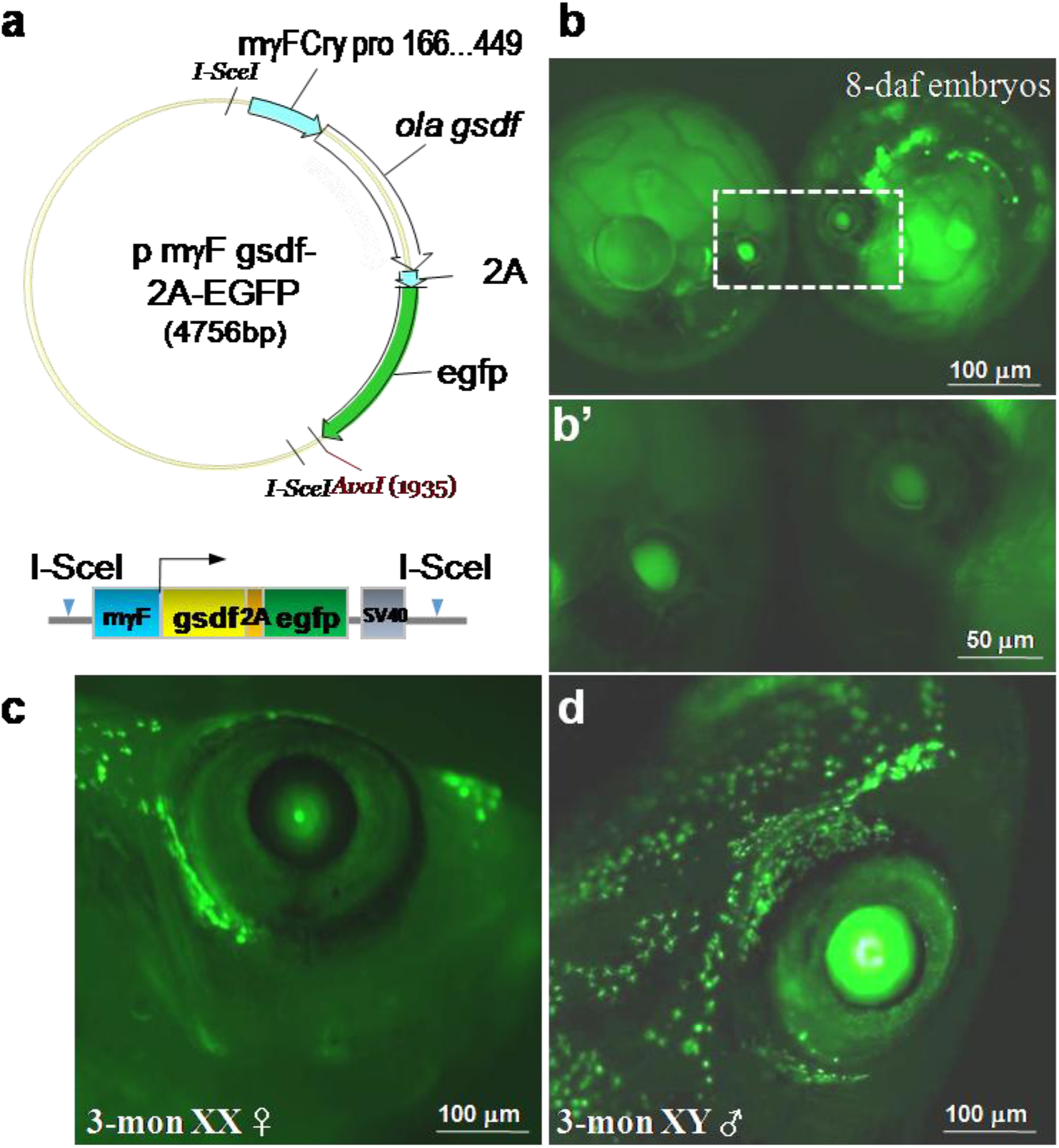
pmFγ-cry reporter driving *gsdf* over-expression in the lens of transgenic medaka. (a) Map and schematics of expression vector. Expression of EGFP was observed in embryos lens from 4 days after fertilization to adulthood. Eight days after fertilization (8-daf) embryos at low magnification (b) and high magnification (b’); 3-month-old adults of F2 progeny (c and d).

GFP signals first appeared in the lens at stages 28-30, right after the onset of retinal pigmentation according to findings by Iwamatsu(Iwamatsu, 2004). The signals became stronger in later embryonic stage (Fig. 1b and 1b’), and were preserved until adulthood (Fig. 1c and 1d). This was consistent with GFP signals from original p817-EGFP reported previously (Vopalensky et al., 2010). Interestingly, GFP signals appeared somehow in a sexually dimorphic fashion, weaker in females than in males (Fig. 1c and 1d), in the progeny of Tg^cryG^ fish from founder 1 male outcrossing to the wild-type XX female, but no difference was observed in offspring from families of founders 2 and 3, nor in the p817-EGFP HdrR strain as a control line (data not shown). This sexual distinction was not reported in previous p817-EGFP Cab and Heino strains(Vopalensky et al., 2010). Whether the distinction was caused by the distinct localization or the number of Tg^cryG^ copies intergrated, coexpression of Gsdf, or a little difference in genetic background between HdrR and Cab/Heino strains needs further investigation.

Extralenticular expression of Gsdf–GFP in transgenic were also found in brain and gonad of both transgenic males and females (Fig. S1). However, no fluorescent signals were detectable in the brains of wild-type males and females as a control (Fig. S1). This extralenticular expression was not described in the transgenic p817-EGFP line(Vopalensky et al., 2010), but was also observed in the p817-EGFP control line (data not shown). Mouse βB2- and γ-crystalline promoters are expressed in the retina, brain, and testis outside lens(Magabo et al., 2000; Templeton et al., 2013), highlighting the possibility of extralenticular expression of gsdf2AEGFP in HPG axis within the transgenic line. This extralenticular expression was detectable in progenies of four subsequent generations used to be investigated.

### XX maleness deduced from Tg^cryG^ fish

Previous studies reported that the over-expression of *gsdf* under the control of β-actin resulted in XX masculinization(Myosho et al., 2012; Zhang et al., 2016). In the pγcryF *gsdf–gfp* transgenic (Tg^cryG^) line, XX males were obtained through four generations of three Tg^cryG^ lines, at the ratio ranging from 17.4% to 44.4% in the progenies of each generation. The breeding data are summarized in Table S2. In the F1 generation, 7 of 32 (21.9%) and 12 of 28 (42.9%) XX males were obtained from 2 independent transgenic lines (Table S2). These transgenic F1 males were further out-crossed to Wt females, generating nearly 50% *gfp*-positive XX males. It is speculated that the mouse γ-crystalline promoter may only drive the expression of *gsdf–gfp* in a few cells not enough to initiate testicular formation in XX Tg^cryG^ female gonads, indicating that the minimum threshold of *gsdf* expressing cells were required for XX maleness. The anatomical analysis was performed to testify this presumption. Surprisingly, two types of GFP signals were observed in transgenic fish: one was XX female with GFP positive in hypothalamus and ovary, but GFP negative in the pituitary region (Figure S1c1-c4), the other was GFP positive in hypothalamus–pituitary–testis (HPT) tightly associated with XX males (Figure S1d1-d4), in agreement with the gonadal expression of *gsdf* causing XX maleness(Myosho et al., 2012; Zhang et al., 2016). In the fourth generation of family 1, 12 of 16 (75%) XX males exhibited GFP-positive signals in HPT; 4 were positive in H but negative in PT. In contrast, 18 of 25 (72%) XX females were gfp positive in H and O, but 7 individuals were GFP undetectable in HPO. Endogenous *gsdf* was detected in the preoptic nucleus of Wt males (Fig. 2a), and weakly detected in few cells co-expressing of cyp19b in the pituitary (Fig. 3a, arrowheads), but undetected in Wt females (data not shown) with ICH analysis. In Tg^cryG^ XX males, however, *gsdf–cyp19b* coexpressing cells or solely gsdf-expressing cells were present in the pituitary (Fig. 2b). The *gsdf-cyp19b* coexpressing cells in the pituitary were distinct from cyp19b-expressing cells in the spinal cord identified in the longitudinal section of whole brain, where they were drastically enriched in XX Tg^cryG^ males (Figure S2).

**Figure 2.**
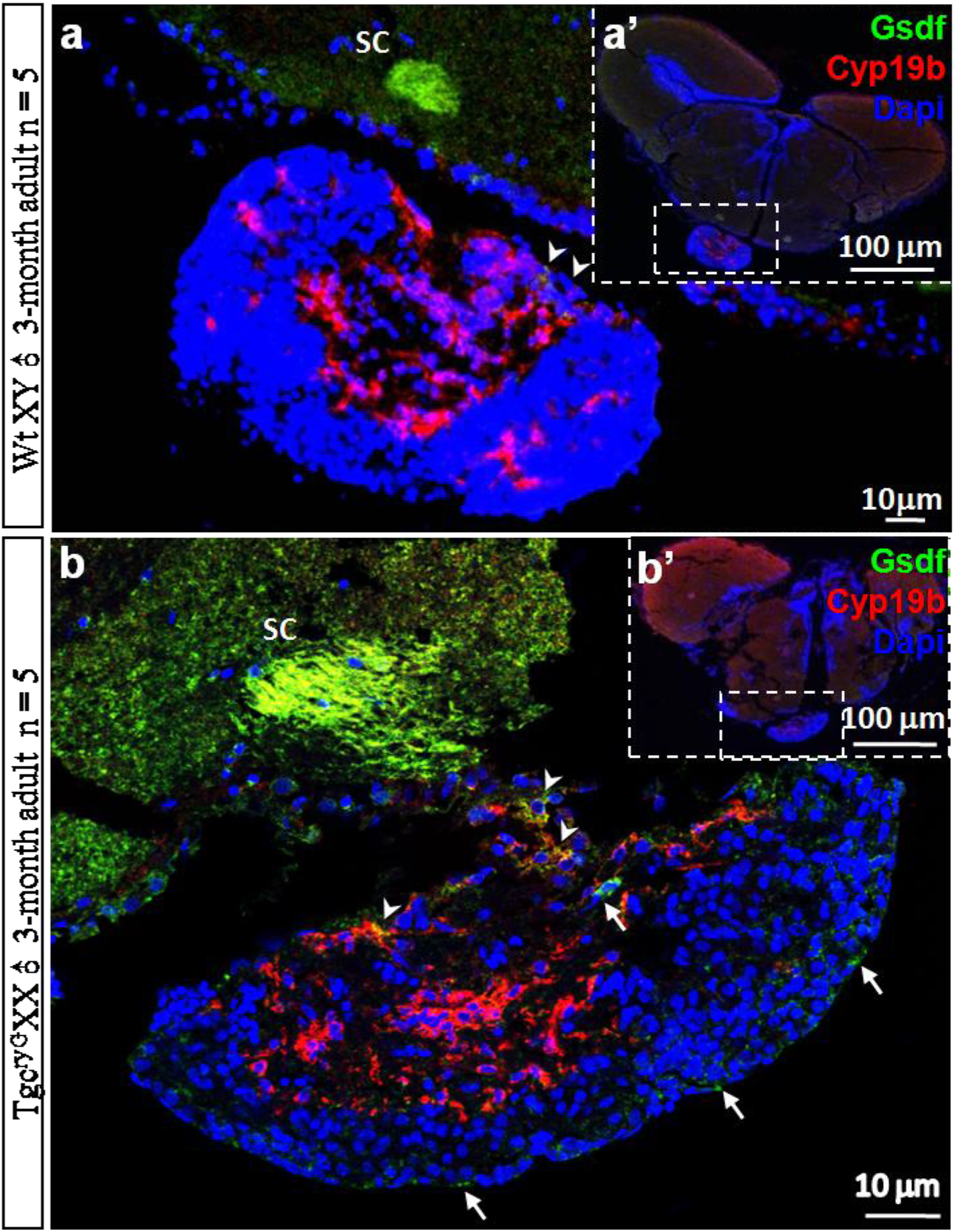
Coexpression of *gsdf* and *cyp19b* cells in Wt XY male and Tg^cryG^ XX male brain. Immunocytohistologic analysis of Wt XY ♂ (a high-magnification image of a’ cross-section); and Tg^cryG^ XX ♂ (b high-magnification image of b’ cross-section) with anti-Cyp19b and Gsdf antibodies. Endogenous Gsdf was detected mainly in suprachiasmatic nucleus (SC) and Cyp19b-positive neurohypophsis cells (arrowheads). Anti-Gsdf reactive signals were found solely (arrows) in the transgenic pituitary.

**Figure 3.**
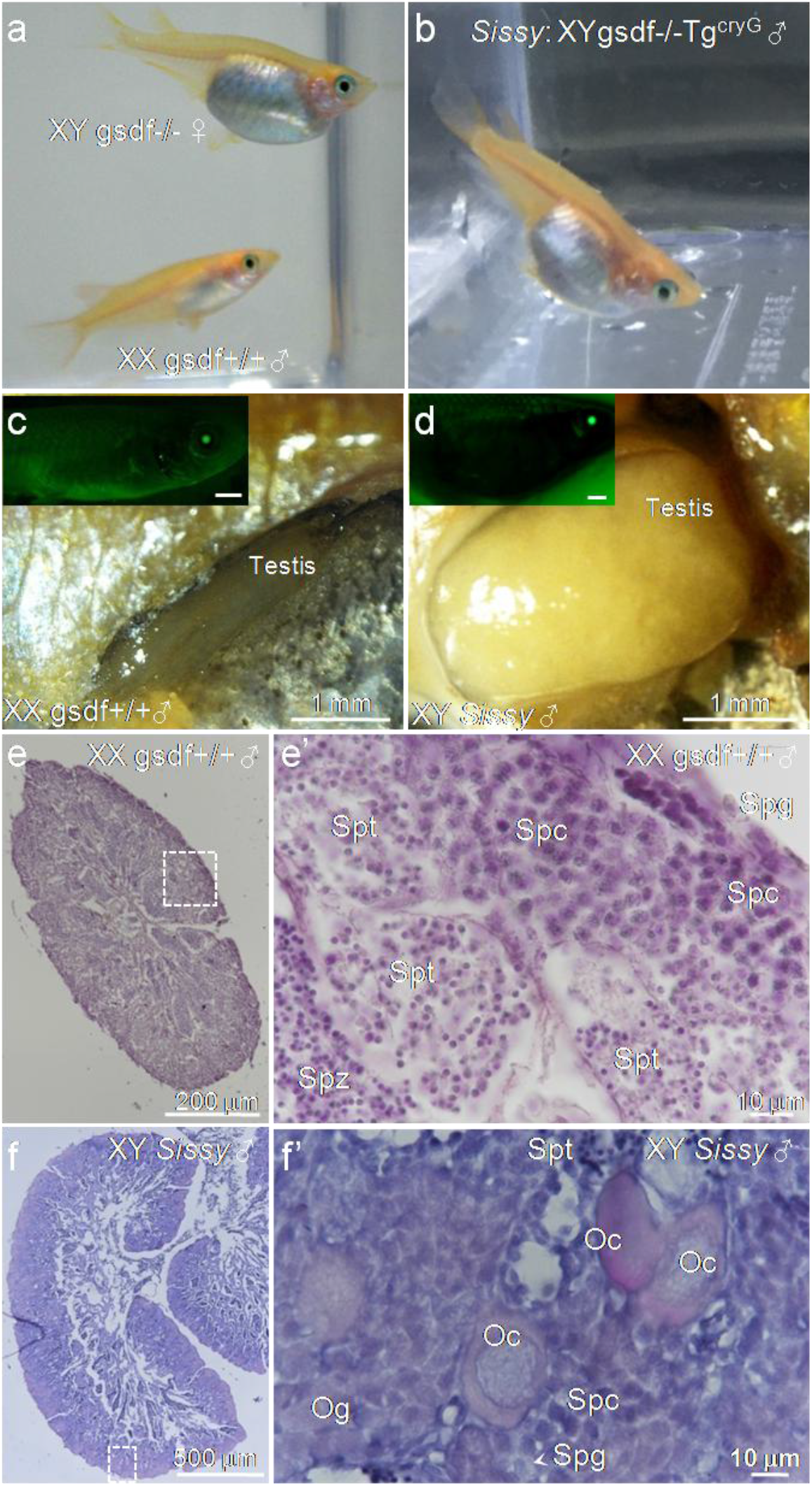
Male development in transgenic lines of *gsdf* intact XX or *gsdf* null XY mutants. (a) Males were obtained from transgenic XX lines, but females from *gsdf* null XY mutants. (b) *Sissy* males were obtained from the XY gsdf null Tg line. Morphology of gonads from XX male (c) and XY *gsdf* null male (d) respectively, with Gfp signals in eyes in upper panels in (c) and (d). (e) HE staining of testis from Tg^cryG^ XX males. (e’) Magnified image of flame in (e). (f) Testis from *Sissy* males. (f’) Magnified image in (f). Spg, spermatogonia (arrowhead); Spc, spermatocyte; Spt, spermatid; Spz, spermatozoa; Og, oogonia; Oc, oocyte.

### Ovotestis development in *Sissy* medaka (Tg^cryG^ *gsdf*^−/−^)

A Tg^cryG^ *gsdf*^−/−^ fish was obtained from the Tg^cryG^ line outcrossing to *gsdf* heterozygotes and subsequent breeding with an XY Tg^cryG^ gsdf^+/-^ male vs an XX gsdf^−/−^ female. The genotypic analysis of *gsdf* and *dmy* is summarized in Table 1. In contrast to six Tg^cryG^-negative XY *gsdfKO* females, as shown in a previous report demonstrating that *gsdf* disruption resulted in XY female sex reversal (Guan, 2017; Zhang et al., 2016), 11 Tg^cryG^ XY *gsdf*^−/−^ males and 8 Tg^cryG^ XX *gsdf*^−/−^ females were obtained from 61 offspring, with a ratio of 1:1 (heterozygotes: homozygotes = 31:30) segregation in the expected Mendelian manner (Table 1). All 11 Tg^cryG^ XY *gsdf*^−/−^ males displayed a pronounced abdominal expansion and a parallelogram-shaped anal fin similar to that of a typical wild-type male (Fig. 3a and 3b). The testis was four-to fivefold bigger than that of a normal XY or XX male (Figure 3c and 3d). HE staining revealed that spermatogenesis proceeded in cysts with variant stages of germ cells from distal spermatogonia to sperms located most proximally (Fig. 3e–e’). However, the distal layer with maximum spermatocytes was thicker than the one in wild-type males (Fig. 3a and 3b). Abundant primary oocytes were scattered adjacent to testicular cysts shown in the magnified image (Fig. 3f–f’), indicating that both oogenesis and spermatogenesis coexisted within one gonad, the gonadal mosaicism observed only in hermaphrodites. No vitellogenic oocytes were found in all 11 ovotestes. The fertilization rate and embryo viability of *Sissy* sperms were evaluates in comparison of wild-type XY sperms with an intact *gsdf* as a control. Results of breeding test were summarized in Table 2.

**Table 1.**
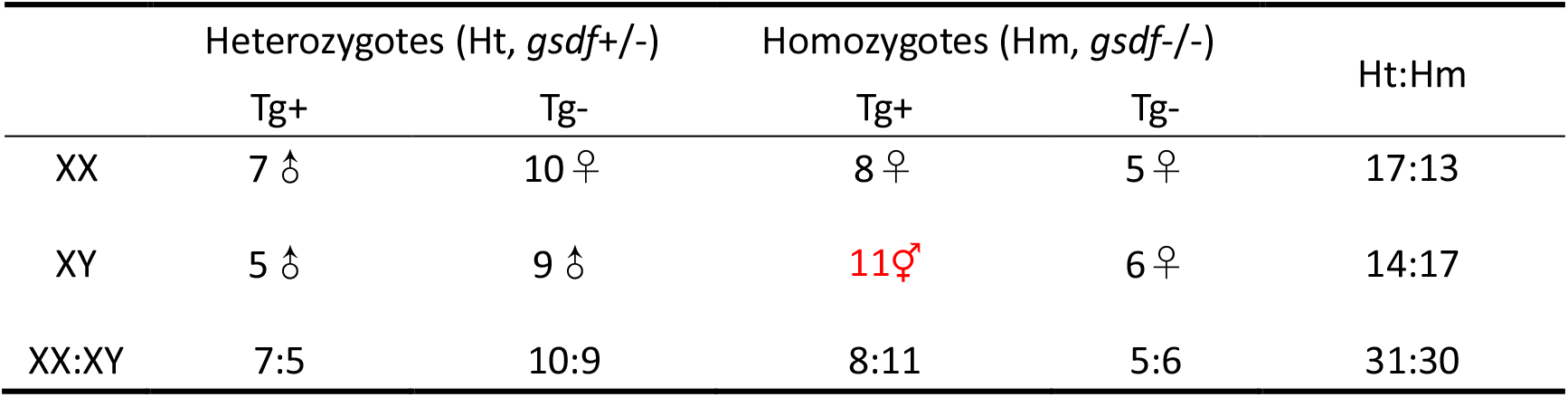
Genotyping analysis of offsprings derived from parental Tg gsdf+/-XY ♂ and gsdf-/-XX ♀

**Table 2.** Sperms from *Sissy* medaka were subfertility revealed by the lower fertilization rate and embryo viability in comparison of wild-type XY sperms with an intact *gsdf* as a control.

|  | Mating | Ovulated eggs | Fertilized eggs (F/O) | Early blastula stage eggs (blastula/F) | Hatching (hatching/F) |
| --- | --- | --- | --- | --- | --- |
| Sissy1♂ | WtXX ♀ | 117 | 53(45.30%) | 49(92.45%) | 45(84.91%) |
| Sissy2♂ | HtXX ♀ | 129 | 33(25.58%) | 26(78.79%) | 23(69.70%) |
| WtXY ♂ | WtXX ♀ | 189 | 189 | 189(100%) | 189(100%) |
| WtXY ♂ | HtXX ♀ | 173 | 173 | 173(100%) | 173(100%) |

### Coexpression of *gsdf2Agfp* and *cyp19b* detected in the pituitary and testis

Despite weak expression of *gsdf* detected in the region of wild-type XY male brain using ICH with monoclonal antibody against Gsdf protein and polyclonal antibody against Cyp19b (Fig. 2), ectopic expression of *gsdf–gfp* was detected in the hypothalamic pituitary region of Tg^cryG^ adult brains, with a large number of *gsdf* and *cyp19b*-coexpressing cells in the pituitary (Fig. S2). In the wild-type XY testis, *gsdf* was detected mainly in *Sertoli* cells surrounding spermatogonia type A and B (Spg A and B) in the most distal region, while the expression of *cyp19b* was high in the proximal region in the epithelium near the efferent duct where mature sperms were released (Figure 4a_1-5_). *Sertoli* cells surrounding Spg B in Tg^cryG^ XX testes were highly reactive to both α-Gsdf and α-Cyp19b antibodies, which was revealed by strong yellow signals from the merged image of green and red fluorescence phases (Fig. 4b_1-5_), indicating that *gsdf*- and *cyp19b-* coexpressing cells increased in Tg^cryG^ XX males compared with wild-type XY testes (Fig. 4a_1-5_). In *Sissy* ovotestes, sole α-Gsdf reactive signals were barely detected in the distal region due to the lack of endogenous *gsdf* expression, comparing to merged signals in orange color surrounding Spg B and spermatocytes (Spc), were positively reacted to both α-Gsdf and α-Cyp19b antibodies. The α-Gsdf-reactive cells were derived from the transgenic *gsdf-gfp* expression, as the endogenous *gsdf*-expressing cells were absent in the ovotestes of *Sissy* males (Fig. 4c_1-5_)(Zhang et al., 2016).

**Figure 4.**
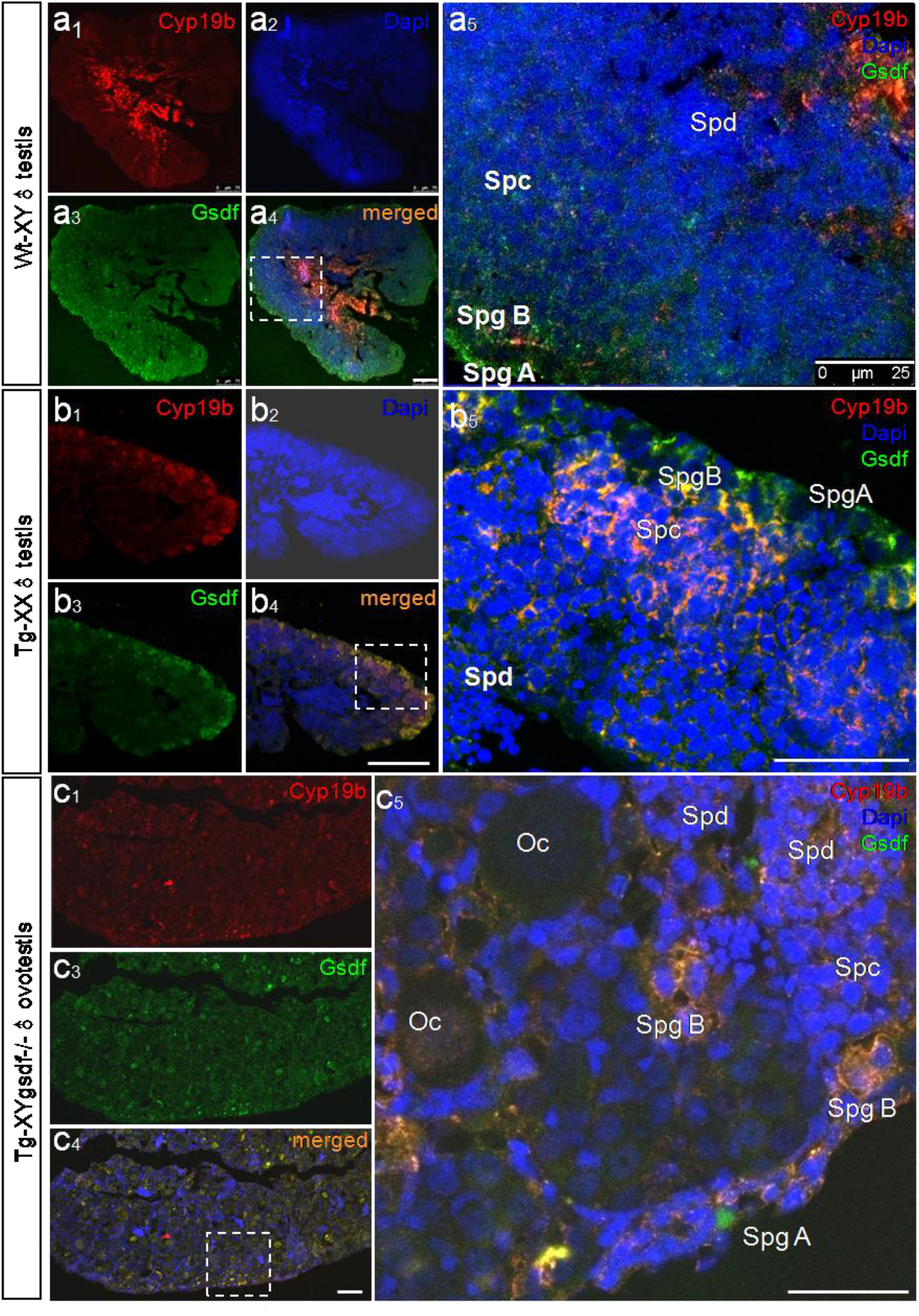
Coexpression of *gsdf* and *cyp19b* increased in Tg adult testis and ovotestis but not in wild-type XY testis. (a1–5) HdrR wild-type adult testis. Gsdf was predominant in Sertoli cells surrounding spermatogonia and *cyp19b* in epithelial cells of efferent duct (a1–4). (a5) Magnified image of a4. (b1–4) Testis derived from an XXTg male with the intact *gsdf* gene. (b5) Magnified image of b4. (c1–4) Testis derived from *Sissy* male with the intact *gsdf* gene. (c5) Magnified image of c4. Sc, Sertoli cells; SpgA & B, spermatogoniaA and B; Spc, spermatocyte; Spz, spermatozoa; sperm. Scale bar: 100 μm in a4, b4 & c4; 25 μm in a5, b5, and c5.

### Mis-*gsdf* signals leading to an alteration of steroidogenesis and estradiol/testosterone (E2/T) balance

The expression of key enzymes responsible for steroidogenesis and serum sexual hormone levels of E2 and T among Wt males, Wt females, Tg^cryG^ XX males, and Tg^cryG^ XY gsdf^−/−^ fish was compared to evaluate the effects of *gsdf* signaling in steroidogenesis. The expression profiles of relevant genes were documented using real-time PCR (Fig. 5 and Fig. S3). The expression of 17α-hydroxylase (*cyp17a*) and *amhr2* was higher in male testes (molecular markers corresponding to testis development), in contrast to *cyp19a*, whose expression was higher in female ovaries (molecular marker corresponding to ovary development). The expression of *cyp19a* expression was inhibited in *gsdf* KO ovaries (Fig. S3A), in agreement with folliculogenesis arrest at the previtellogenic stage in *gsdf* deficiency(Guan, 2017). The expression of *cyp19a* was also lower in Tg^cryG^ XX testis and Tg^cryG^ *gsdf*^−/−^ ovotestis as expected (Fig. S3A).

**Figure 5.**
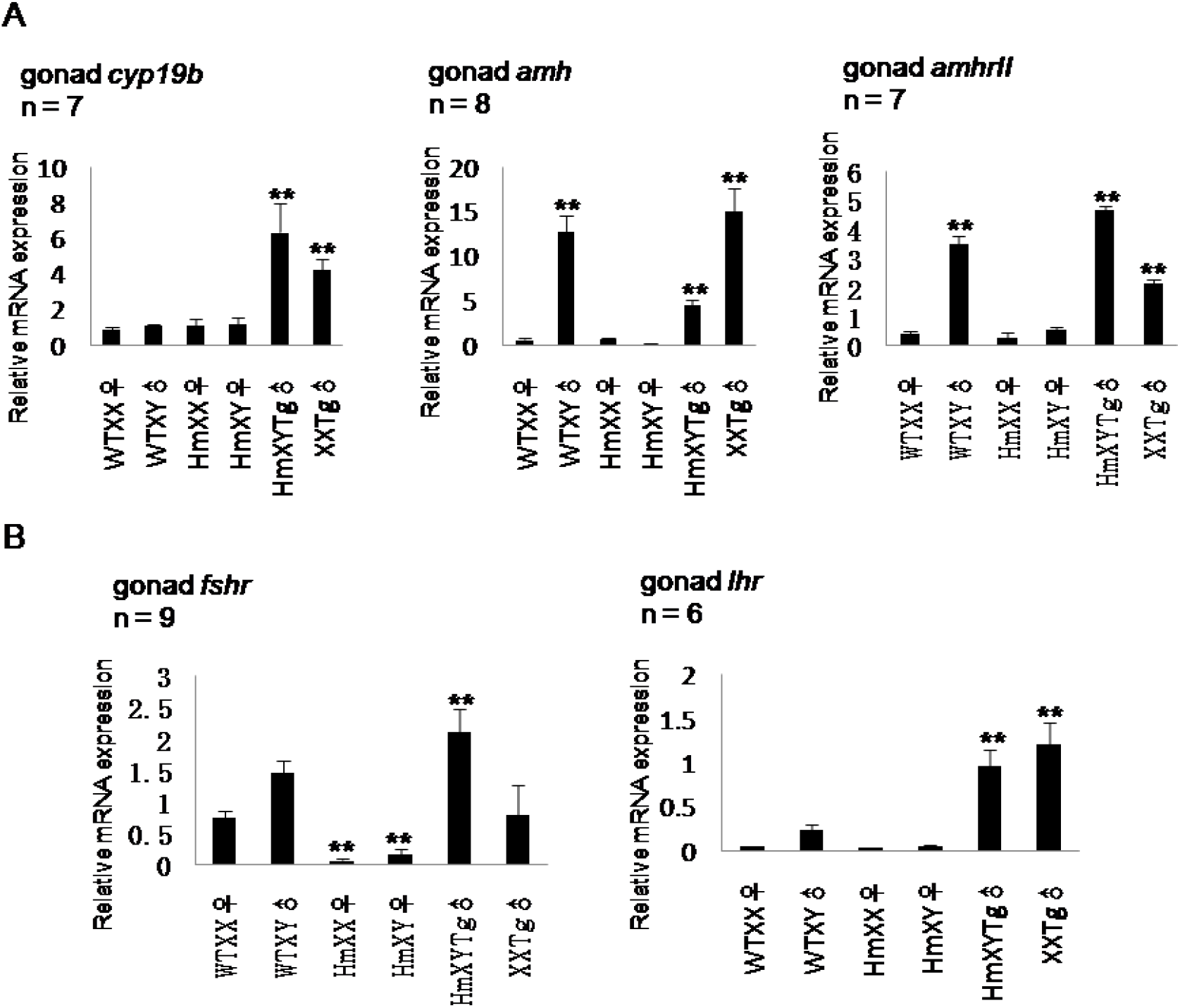
Transcript response profile of *gsdf* deficiency or transgenic *gsdf*. (A) Brain-type *cyp19b* had no sexual dimorphism between males and females, but was greatly induced in transgenic XX testes and *gsdf* KO ovotestes. Also, *amh* was higher in male testes than in female ovaries. The same pattern was revealed by its receptor *amhr2*. However, a drastic increase in *gsdf* KO Tg ovotestes was observed, likely due to the super-response to exogenous *gsdf* stimulation in *gsdf* KO ovotestes. The absence of *gsdf* decreased *amh* expression in ovotestis, but increased *amhr2* expression, indicating a potential *gsdf–amhr2* binding. (B) Transgenic *gsdf* elevated the expression of *fshr* and *lhr*, while the loss of *gsdf* inhibited the expression of *fshr* and *lhr*. ** *P* < 0.01.

Serum levels of E2 and T were evaluated using an enzyme immunoassay kit previously described(Chen et al., 2014). The results of E2/T assay are summarized in Figure S3B. The absolute serum value of E2 was higher than that of T in wild-type females, but was lower in wild-type males, *gsdf*^−/−^ females, Tg^cryG^ XX males, or *Sissy* males. The ratio of E2 vs T was almost 100 times higher in Wt females than in Wt males (Fig. S3B), resulting from the effective *cyp19a* expression higher in ovary than in testis. Plasma E2 in *gsdf*^-/^ mutants was one fourth of that in normal females, and no obvious T alteration consistent with the low expression of *cyp19a* in *gsdf* depletion. In contrast to a low level of E2 unchanged in transgenic lines, the testosterone level was elevated in transgenic XX males and *Sissy* lines, suggesting T synthesis was greatly stimulated by *gsdf* transgenic signals (Fig. S3B), leading to a decrease in the E2/T ratio to a normal male level and the fin shape resembling that of a normal male in XX transgenic males. The E2/T ratio of *Sissy* was one half to one fourth of that in *gsdf* KO females and almost one eighth of that in normal females, explaining male sexual characterization externally in the form of anal fin shape but ovotestis formation internally (Fig. S3B).

The expression profiles of *cyp19b, amh/ahmrII*, and *fsh/lh* receptors were assessed using qPCR analysis and compared among *gsdf* deficiency, *gsdf* transgenic lines, and normal fish as a control to investigate the consequence of E2/T and gonadotropin feedback in HPG axis. *cyp19b* showed no sexually dimorphic expression in males and females; however, the expression increased drastically in transgenic lines (Fig. 5A).

Both *amh* and *amhr2* were higher in the Wt adult testis than in the Wt adult ovary, consistent with a previous observation(Kluver et al., 2007). They were inhibited on *gsdf* depletion but increased in transgenic *gsdf* lines, suggesting the expression sensitivity to transgenic *gsdf* signals (Fig. 5A). Both *fshr* and *lhr* decreased in *gsdf*^−/−^, consistent with a previous report (Fig. 5B) (Kaiqing Sun1#, 2017). Interestedly, *fshr* and *lhr* greatly increased in *Sissy* fish, indicating that transgenic *gsdf* could stimulate their expression probably via paracrine or endocrine effects (Fig. 5B).

### Decrease of α-Lh and α-Cyp19b cells in pituitaries in transgenic *Sissy* lines

Lh-secreting cells were severely inhibited in *Sissy* pituitaries compared with those of Wt males (Fig. 6). Some Lh-positive cells localized near Cyp19b-positive cells but did not overlap, indicating that transgenic *gsdf* signals from cyp19b-positive cells in Tg^cryG^ lines has a paracrine effect on Lh-secreting cells probably upon a gradient of *gsdf* secretion. In gonads, α-Cyp19b and α-Lh reactive cells were not overlapped but adjacent to each other (own data unpublished).

**Figure 6.**
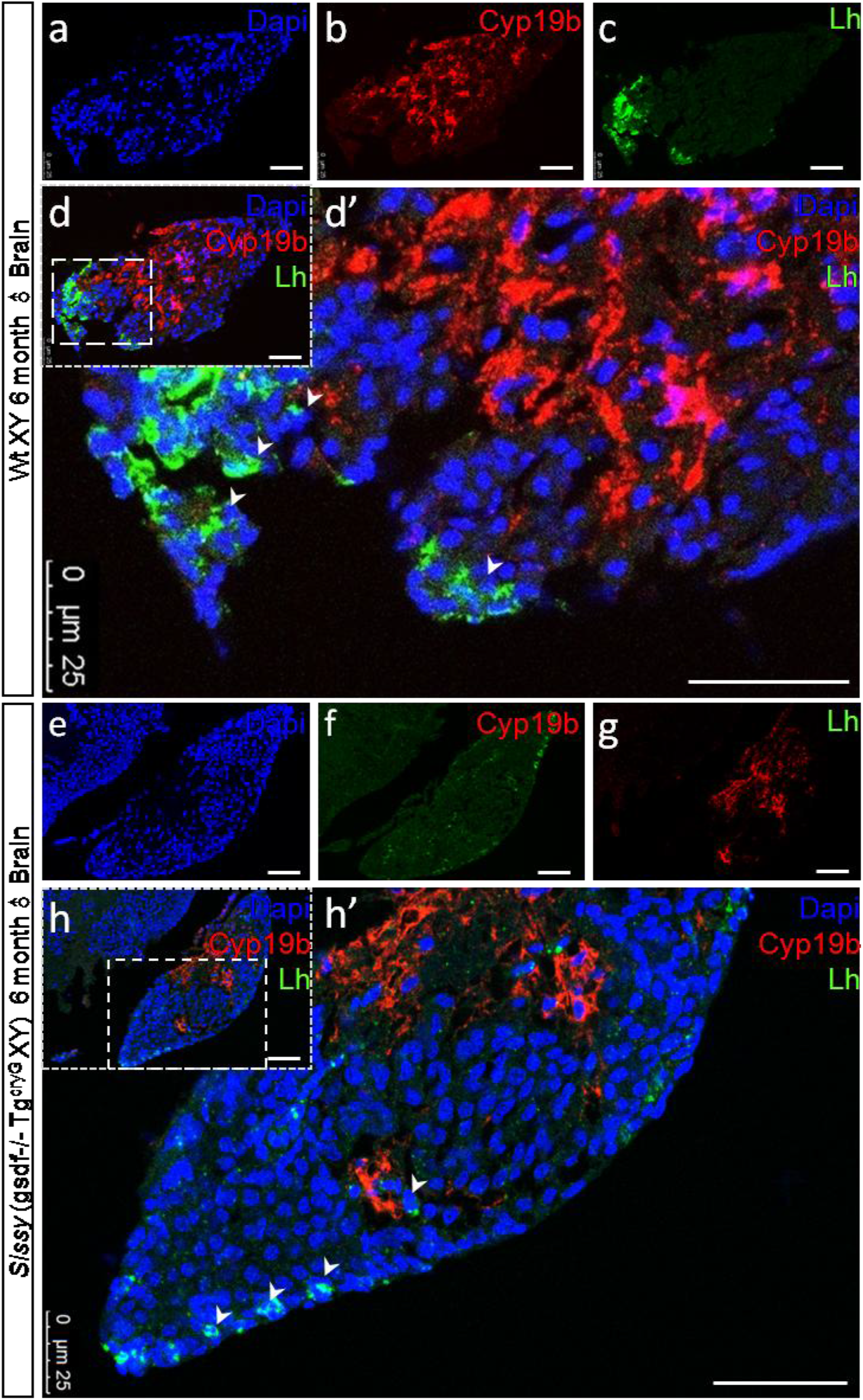
Cyp19b and Lh inhibition in *Sissy* pituitary. (a–d) ICH result of Wt♂ pituitary with anti-Cyp19b & anti-Lh. (e–h) *Sissy♂* pituitary. Arrowheads represent Lh-positive cells. Scale bar: 50 μm in a-d and e-h; 25 μm in d’ and h’.

### Hypothesis regarding the role of *gsdf* in gonadal differentiation

Based on the aforementioned data, a hypothesis was formulated regarding the role of *gsdf* in gonadal differentiation (Fig. 7). *gsdf* probably initiates male development by coupling with *amh* and binding with *amhr2* and/or *gsdf-specific* receptor. *cyp19b, fshr* and *lhr* are downstream genes controlled by *gsdf* signals. Gsdf disruption reduced fertility and fecundity by affecting HPG gonadotropins and controlling the release of sex steroids. Blocakge of *amh/gsdf* signaling resulted in ovotestis formation with both spermatogenesis and oogenesis coexisting in one gonad in medaka.

**Figure 7.**
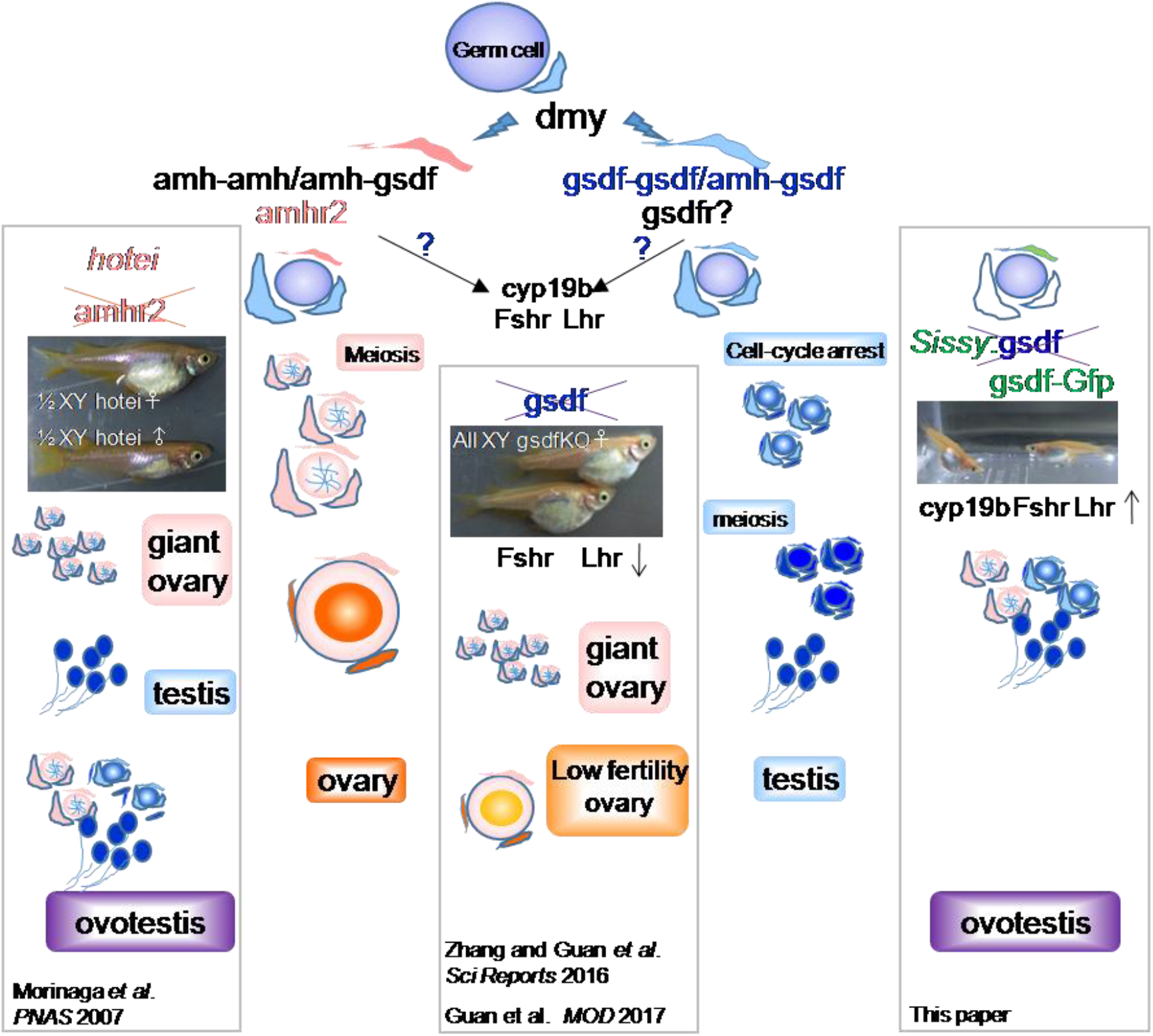
Schematic diagram of gonadal differentiation under *gsdf* control. Gsdf probably initiates male development by coupling with *amh* and binding with *amhr2* and/or -specific receptor. *cyp19b, fshr*, and *lhr* are downstream genes controlled by *gsdf* signals. *gsdf* disruption reduced fertility and fecundity by affecting HPG gonadotropins and controlling the release of sex steroids. Blocakge of *gsdf/amh/amhr2* signaling resulted in ovotestis formation with both spermatogenesis and oogenesis coexisting in one gonad in medaka.

## Discussion

### Similarities in organization of HPG and lens between fish and mammals

This study showed that the mouse γ–crystallin promoter was capable of directing the production of Gsdf–Gfp fusion proteins in medaka lens, brain and gonad, indicating the common *cis*- and *trans*-element sharing between fish and mammals responsible for the organization of HPG axis and lens during early embryogenesis. HPG axis control of gonadal development, as well as sexual behavior, is generally conserved in all vertebrates. Despite the morphological difference present in the pituitary gland between fish and humans, the circulation of gonadotrophins from the pituitary and functions by binding to their specific receptors in controlling the process of folliculogenesis and spermatogenesis is well conserved from fish to mammals(Murozumi et al., 2014). Therefore, *Sissy* medaka enables *in vivo* and *in vitro* analysis of molecular events under HPG gsdf/TGF-β signaling regulation of gonadal differentiation.

### *gsdf/amh* inhibition of germ cell proliferation is essential for spermatogenesis

Although sex determining system is highly diverse in vertebrates, it is common that the primordial germ cells (PGCs) reach the gonads in female to undergo cell proliferation and oogenesis, in contrast to the retaining of mitotically quiescent state for a while before the initiation of spermatogenesis in male gonads. Indifferent germ cells require instructive signal(s) from the surrounding somatic cells to enter their sexual path, in other words, the initiation of oogenesis and spermatogenesis controlled by the somatic cells steps by steps(Tanaka, 2016). Genetic manipulation of germ cell *foxl3* gene (*forkhead*-domain gene 3 as an intrinsic determiner of sperm–egg fate) or the germ cell numbers in medaka disrupt the communication between germ cells and surrounding somatic cells, and subsequently result in ovotestis development(Tanaka, 2016). *foxl3* expression was merely changed among the adult testes of normal XY males, transgenic XX males and *Sissy* intersexes (own data unpublished), suggesting a rare alteration of germ stem cells under the *gsdf* signals influence. Although the expression of *cyp19a* reduced in *gsdfKO*, a slight increase in transgenic lines compared with normal XY males, no apparent change between removing or adding *gsdf* were found, suggesting that the decrease in the expression of *cyp19a* may be an indirect consequence of *gsdf* deficiency, probably due to the halting of oocyte development. Ovotestis developed in the adults of *cyp19aKO* could be under the influence of a pathway from the one initiated by gsdf(Nakamoto et al., 2017). In contrast to *cyp19a, cyp19b*, as well as *amh* and *amhr2*, was greatly induced in transgenic *gsdf* lines. High expression of *amh* in pre-Sertoli cells is reduced with the onset of androgen activity at puberty in mammals, birds and reptiles, in contrast to the persistent *amh* expression until adulthood in fish probably due to the cystic structure of fish testis(Pfennig et al., 2015). Lack of *gsdf* signaling or *amh* signaling via *amhr2*, directly or indirectly leads to sex reversal from male to female(Imai et al., 2015; Morinaga et al., 2007; Zhang et al., 2016). A remarkable similarity between *gsdf* null mutants and *hotei* phenotypes, including an excessive proliferation of germ cells and oocyte arrest at the previtellogenic stage, suggested that *gsdf* probably cooperated with *amh/amhr2* signaling system during gonadal development. Dose-dependent proliferation of type-A spermatogonia was demonstrated under trout *gsdf* signaling *in vitro* testicular cells experiment(Sawatari et al., 2007). Transgenic *gsdf* signaling could stimulate the male-fate spermatogonia A proliferation and resulted in XX masculinization in medaka. The proliferated type A spermatogonia were inhibited by *amh* for further proliferation indirectly via *amhr2* signaling. Therefore, the expression of *gsdf* but not *amh* displays sexual dimorphism during gonadal differentiation in medaka(Kluver et al., 2007; Shibata et al., 2010). It was speculated that transgenic *gsdf* signals enabled spermatogenesis in genetic females, and *Sissy* intersex individuals with Tg^cryG^ XY *gsdf* depletion, was the result of promotion of type A spermatogonia proliferation from the pool of germline stem cells under the system of *gsdf/amh/amhr2*, and/or other TGF-β signalings properly, with intact *gsdf* in XX males but improperly null *gsdf* in *Sissy* individuals. This also explained the scenario occurring in some *hotei* mutants (Morinaga et al., 2007). Miss-signaling in the above system led to impairment of the delicate balance between germ cells proliferation and differentiation, and subsequent ovotestis development endorsed by normal *gsdf* but mutant *amhr2* in these *hotei* mutants. The protein interaction between Gsdf and Amhr2 or other candidates is under investigation using the yeast two-hybrid and dual-membrane system.

### Cyp19b is the key player in HPG axis under *gsdf* signals during spermatogenesis

*cyp19b* is expressed mostly in radial glial cells in the embryonic brain of all vertebrates, disappears at birth in the mammalian brain, but remains abundant in the brain of adult fish, amphibians, birds, and reptiles(Coumailleau et al., 2015). The expression of *cyp19b* in medaka begins just before hatching, coincides with the increase germ cells in female than male embryos(Patil and Gunasekera, 2008). No apparent differences in the expression of *cyp19b* have been observed between males and females in zebrafish(Kallivretaki et al., 2007), but a sexually dimorphic expression after puberty has been found in medaka brain(Okubo et al., 2011), indicating that *cyp19b* is involved in HPG regulation of gonadal development at the onset of spermatogenesis. The expression of both *cyp19b* and *lh* decreased in *Sissy* pituitaries, in agreement with the findings in mice aromatase knockout (ArKO), indicating that aromatase is involved in maintaining the Lh-positive cells in medaka and mouse pituitary(Carretero et al., 2016).

Higher gonada-somatic index (GSI) was observed in *gsdfKO* females(Kaiqing Sun1#, 2017) and *Sissy* intersexes associated with elevation of Fsh in pituitary, supporting that the gonad enlargement resulted from Fsh elevation(Kaiqing Sun1#, 2017) (own data unpublished). An increase in GSI has been observed in *lhKO* medaka(Takahashi et al., 2016), *lhKO* and *lhrKO* zebrafish(Chu et al., 2014), in agreement with findings of less Lh secretory cells and only fshr were elevated in *Sissy* pituitary and gonad respectively, as the result of less Lh but elevated Fsh in *Sissy* pituitary (own data unpublished). In the gonadotrophin-dependent phase of follicular development, androgen was produced by *cyp19b*- and *lh*-positive theca-interstitial cell layer and diffused into the mural granulosa cell layer, which produced E2 with Fsh-sensitive aromatase catalysis. E2/T feedback inhibits pituitary secretion of Fsh, which in turn results in the concentration of Fsh to drop below the threshold level in the developing cohort follicles, thereby arresting the development of Fsh-insensitive follicles(Durlinger et al., 1999). Similar to mammals, the major actions of gonadotropins in teleosts were mediated through steroid production. Fsh and E2 levels simultaneously increased during folliculogenesis, and Fsh stimulated E2 production and altered ovarian aromatase activity(Montserrat et al., 2004). Expression of *fshr* and *lhr* drastically decreased in *gsdf* deficiency, but could be stimulated via transgenic *gsdf* addition, indicating that both *fshr* and *lhr* expressing cells are sensitive to *gsdf* signaling in medaka. Multiple knockout or conditional knockout medaka will be needed to uncover the network and the interactions of these factors essential for germ cell differentiation and conserved development from fish to mammals.

In summary, being a lower vertebrate, medaka is helpful in revealing missing information on gonadal differentiation regulated by HPG axis from the evolutionary viewpoint. A *Sissy* transgenic medaka line was established in this study, which enabled the initiation of spermatogenesis and male development through promoting the expression of *gsdf-cyp19b* in gonad and reducing the expression of *cyp19b-lh* in the pituitary. This model may be helpful in further studies on paracrine and endocrine effects of *gsdf/amh/amhr2* TGF-β signals *in vivo*, whereas a subfertility in both male and female mice has been reported harboring a mutation in βB2-crystallin mutation(Duprey et al., 2007).

## Materials and Methods

### Animals

Medaka fish were housed in re-circulating systems at 26–28 °C, with a light-dark cycle of 14-h daylight and 10-h darkness. They were handled in strict compliance with the guidance of the Committee for Laboratory Animal Research at Shanghai Ocean University. Gsdf disruption was accomplished using zinc finger nuclease previously reported in the HdrR strain(Zhang et al., 2016). Phenotypic sex was assessed using secondary sex characteristics (dorsal and anal fin shape) and confirmed by gonadal biopsy of a 6-month adult fish. Genotypic sex (XX or XY) was assessed using PCR primer sets of PG17.5-17.6 to amplify a dmy-specific fragment and a fragment from *dmrt1* of medaka(Matsuda et al., 2002). Genotyping of the *gsdf* allele was performed with primers as described previously(Zhang et al., 2016).

### Designing of transgenic construct and generation of mγF-cry-gsdf2AEGFP transgenic medaka

Original plasmid pmγF-cry-EGFP was constructed by Dr. Zbynek Kozmik (p817-EGFP, Addgene Plasmid 23156), which contained a 271-bp mouse mγF-cry promoter driving EGFP reporter specifically in the lens, facilitating a preselection of transgenic medaka(Vopalensky et al., 2010). The region of EGFP was replaced by the fragment of gsdf2AEGFP (p817-gsdf2AEGFP), which was effective in XX masculinization via the ubiquitous expression under the control of medaka β-actin promoter(Zhang et al., 2016). Further, 40 ng/μL of testing plasmid DNA (p817-gsdf2AEGFP) or control p817-EGFP was used for pronuclear microinjection into fertilized eggs of HdrR strain at the one-cell stage. The cells expressed EGFP and Gsdf simultaneously due to the coexistence of gsdf2AEGFP. Transgenic lines were established via lens green fluorescence protein (GFP) signals in a progeny founder fish crossing to wild-type fish for germline transmission. Fluorescent signals of GFP were captured with a Nikon Ds-Ri2 system under SMZ stereomicroscope (Nikon, Japan) or photographed with confocal microscopy (Leica DMi8, TCS SP8, Germany).

### RNA isolation, reverse transcription, and real-time PCR

Total RNA was extracted from gonads and reverse transcribed into first-strand cDNA with AMV reverse transcriptase (Takara, Dalian, China) and oligo-dT(_18_) primers according to the manufacturer’s protocol. The primers used for qPCR are listed in Table S3. qPCR was carried out using cDNAs as templates and SYBR Fast qPCR mix (Takara), and detected with 7500 real-time PCR system (Applied Biosystems, CA, USA) to measure fluorescence. Thermal cycling conditions were 1 cycle of 2 min at 95°C followed by 40 cycles of 15-s denaturation at 95°C and a 34-s extension at 64°C. The relative abundance of each sample gene was evaluated using the formula: R=2^-ΔΔCt^. β-Actin was used to normalize expression values. Statistical comparisons were evaluated using analysis of variance followed by a pair-wise Student ttest (*P* <0.05 was considered to be statistically significant unless specified). Data were means of three to five independent experiments.

### Histology, immunohistochemical analysis, and *in situ* hybridization

The gonads were fixed in Bouin’s solution for histological and immunohistochemical analysis, or 4% paraformaldehyde for *in situ* hybridization (ISH). The sections were cut at 5 μm thickness and stained with hematoxylin–eosin (HE) according to standard protocols. ISH was performed with digoxigenin (DIG)-labeled or fluorescein isothiocyanate (FITC)-labeled-UTP under the action of T7 or Sp6 RNA polymerase as described previously (Roche, Germany) (Zhu Yefei, 2016). Hybridization signals were analyzed using alkaline phosphatase–conjugated anti-DIG antibody (Roche, Germany) and further incubated with horseradish peroxidase (HRP)-conjugated anti-DIG or anti-FITC antibodies (MBL, Nagoya, Japan). Immunostaining and nuclear staining of cryosections of gonad and brain using DAPI were performed with mouse monoclonal anti-Gsdf(Zhang et al., 2016) and rabbit anti-GFP (MBL, Nagoya, Japan). Rabbit polyclonal anti-Fsh was a gift from Dr Akio Shimizu according to a previous description(Kaiqing Sun1#, 2017; Shimizu et al., 2012). Information of polyclonal antibodies against medaka Cyp19b and Lh β-subunit, monoclonal antibody against medaka Gsdf were listed in Table S4. Multicolor signals were amplified with a TSA Plus Fluorescein/TMR system (PerkinElmer Inc., MA, USA) according to previously described protocols(Zhang et al., 2016). The signals were photographed with a confocal laser scanning microscope (Leica DMi8 TCS SP8, Germany).

### Measurement of serum estradiol and testosterone levels

Blood samples were collected via decapitated carotid artery of each individual fish using a heparinized micropipette tip. Plasma estradiol (E2) and testosterone (T) levels were measured with an E2 and T enzyme immunoassay kit (Cayman Chemical Company, MI, USA) according to the manufacturer’s protocol(Zhang et al., 2008). Statistical comparisons were evaluated using a pair-wise Student *t* test (*P* <0.05 was considered to be statistically significant unless specified. Data are means ± standard errors of three to five independent experiments.

## ACKNOWLEDGMENTS

The authors are grateful for the gift of anti-Fshβ from Prof Akio Shimizu (National Research Institute of Fisheries Science, Japan). We also thank MedSci (<u>www.medsci.com</u>) for the linguistic assistance during the preparation of this manuscript.

---

**Table S1.**
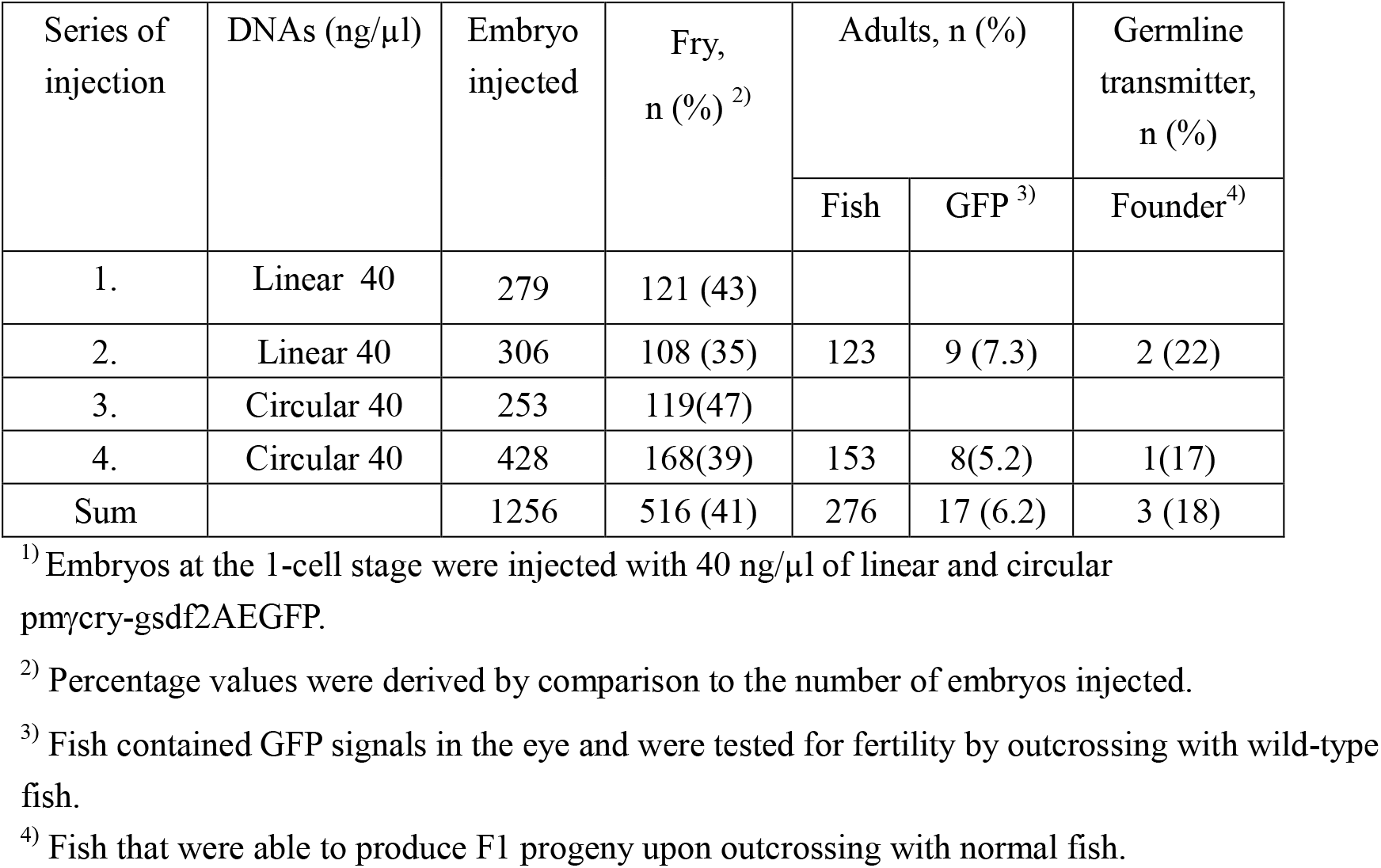
Survival and transgenic efficiency in p817-gsdf2A-EGFP medaka

**Table S2.** XX males development in Tg medaka

| F0 | Family 1<br>Founder 1 XY♂<br>Number of GFP+XX<br>(GFP+XX♂)<br>% | Family 2<br>Founder2 XX♀<br>Number of<br>GFP+XX<br>(GFP+XX♂)<br>% | Family 3<br>Founder2 XX♀<br>Number of<br>GFP+XX<br>(GFP+XX♂)<br>% | Control<br>Wt XX♀<br>Number of<br>GFP+XX<br>(GFP+XX♂)<br>% |
| --- | --- | --- | --- | --- |
| F1 | 32 (7)<br>21.9% | 28(12)<br>42.9% | 23(4)<br>17.4% | 85(0)<br>0% |
|  | F1 XX♂- XX♀ | F1 XX♂- XX♀ | ND | ND |
| F2 | 53(20)<br>37.7% | 20(8)<br>40% | ND | ND |
| F3 | 41(16)<br>39% | 11(39)<br>28.3% | ND | ND |
| F4 | 45(20)<br>44.4% | ND | ND | ND |

ND stands for “Not Done”.

**Table S3.**
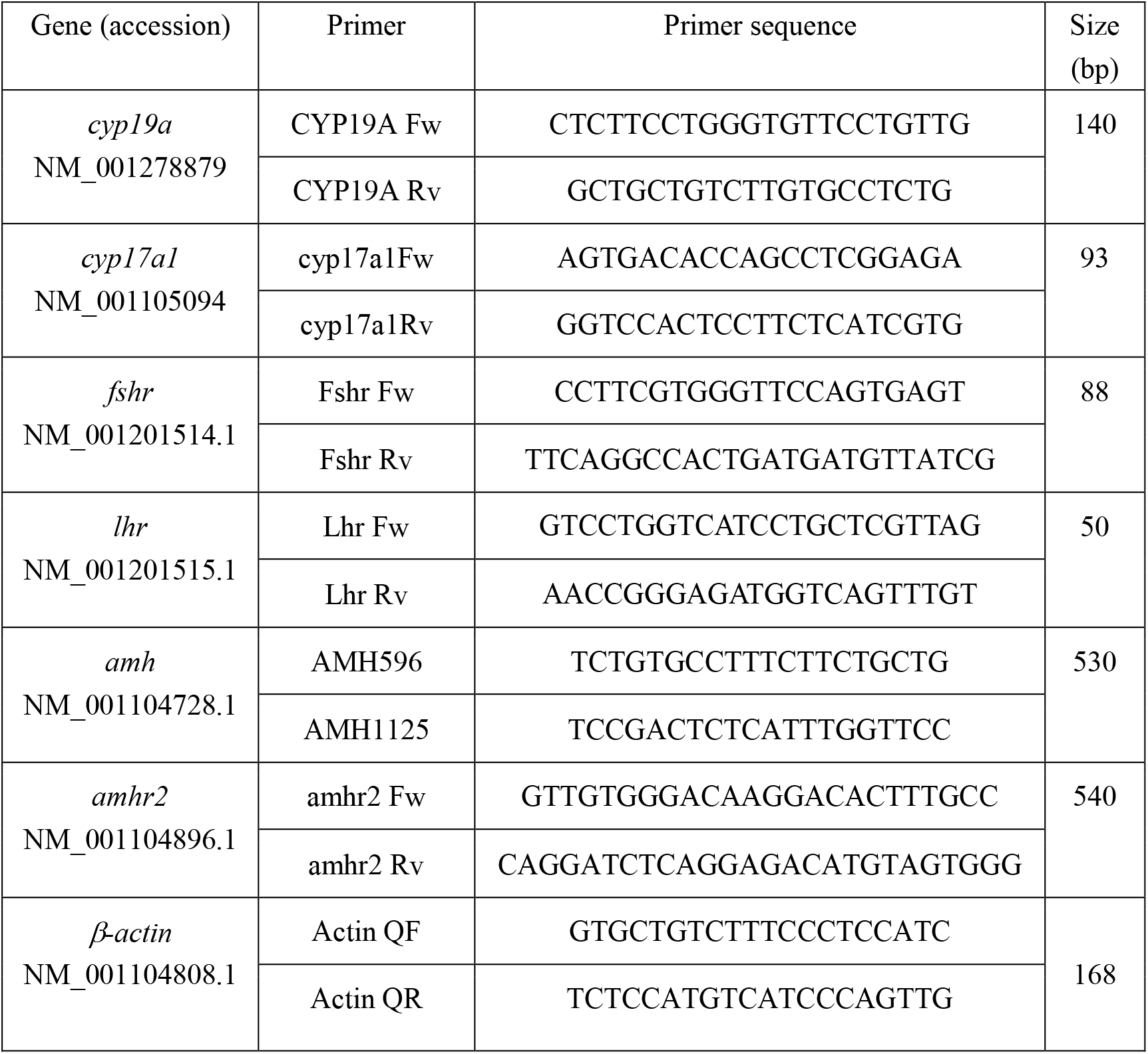
primers for qPCR

**Table S4.**
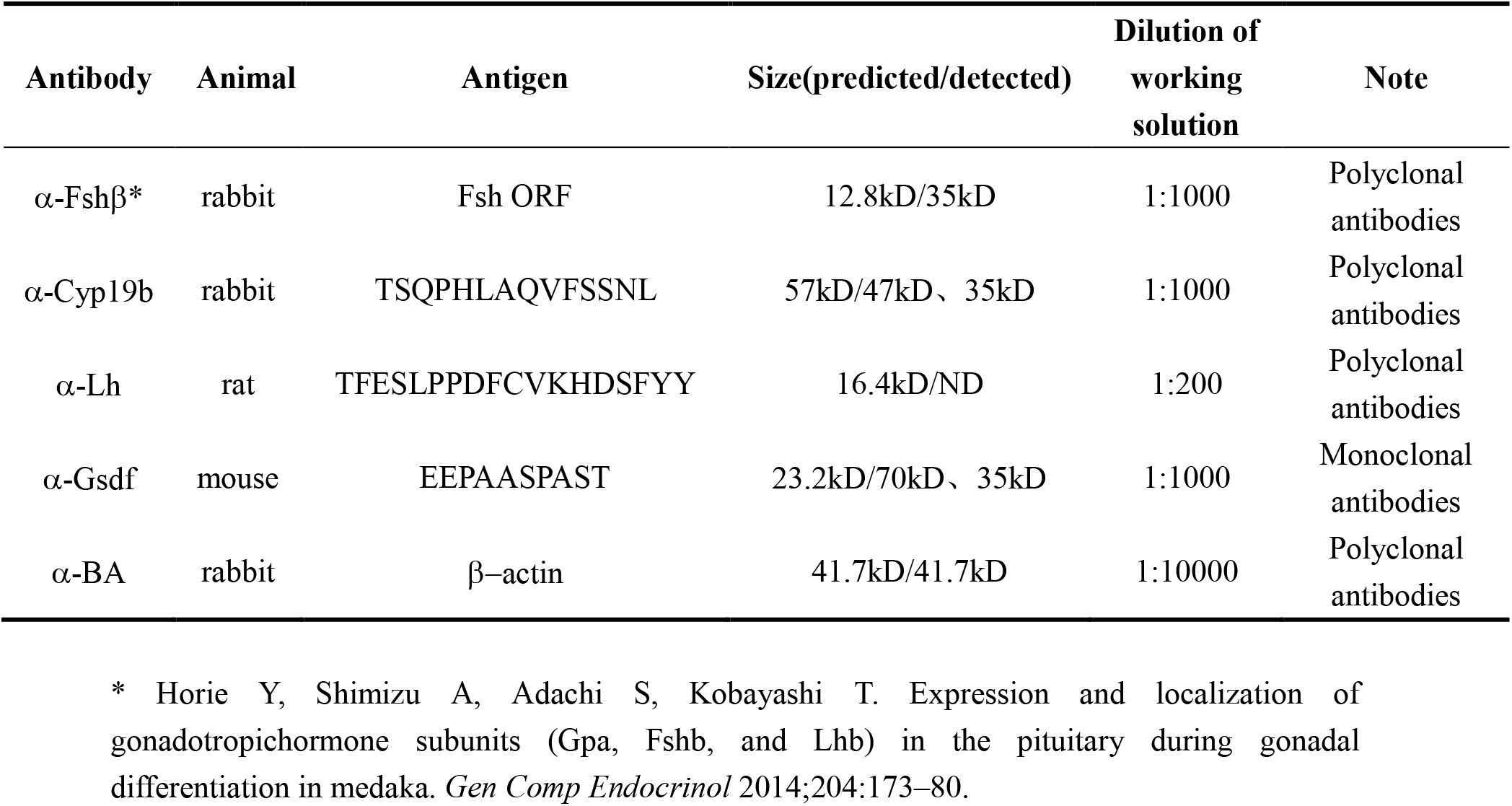
- Results of Western blot analysis using specific antibodies

**Figure S1.**
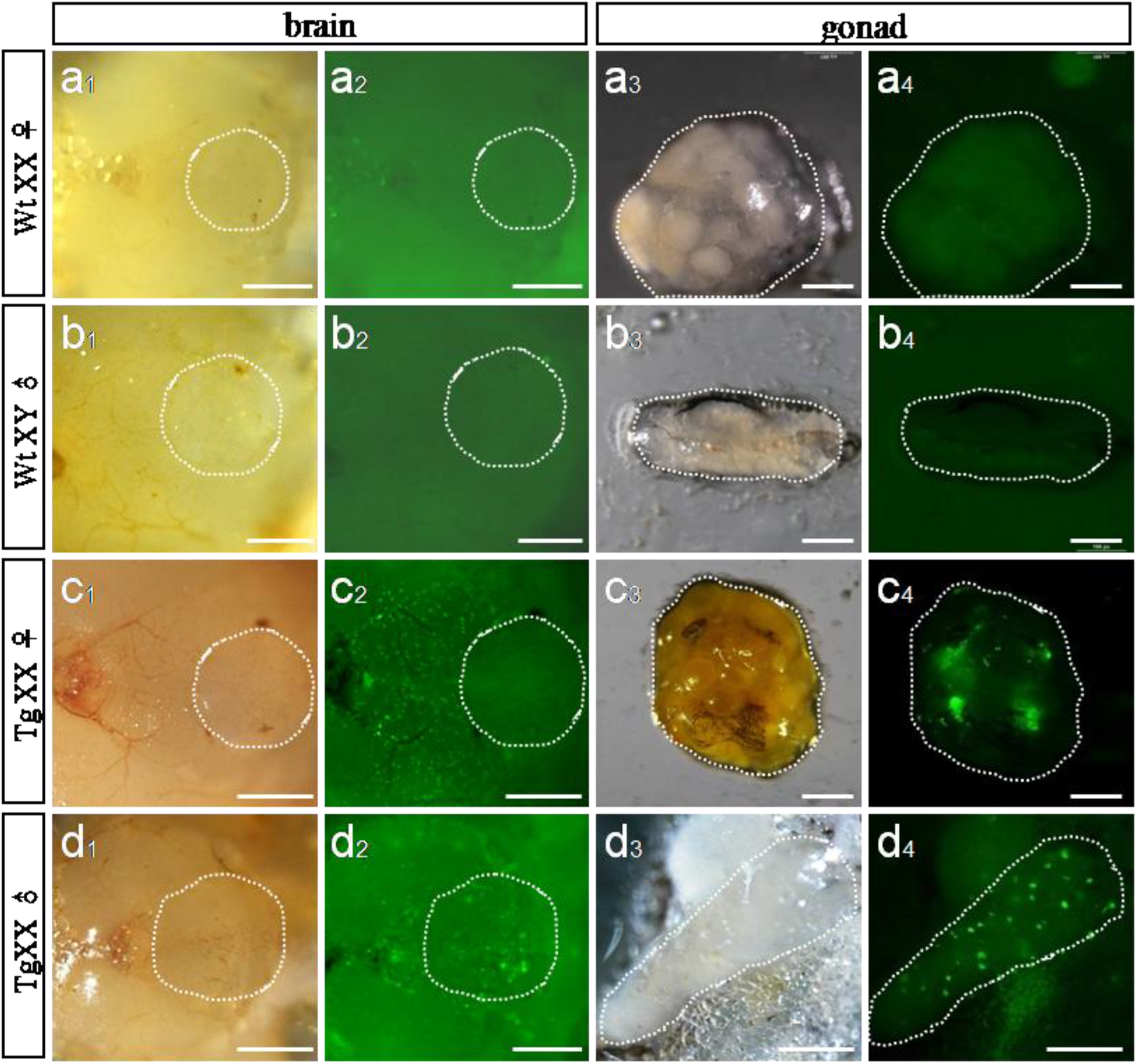
Extralenticular expression of pmFγ-cry-gsdf2AEGFP in brain and gonad. No GFP signals were observed in wild-type XX♀ brain and ovary (a1-4); XY ♂brain and testis (b1-4). GFP signals were detected in 3-month-old transgenic adults of F2 XX ♀ in the brain (c1 and c2) and ovary (c3 and c4); XX♂ brain (d1 and d2) and testis (d3 and d4). Scale bar: 100 μm.

**Figure S2.**
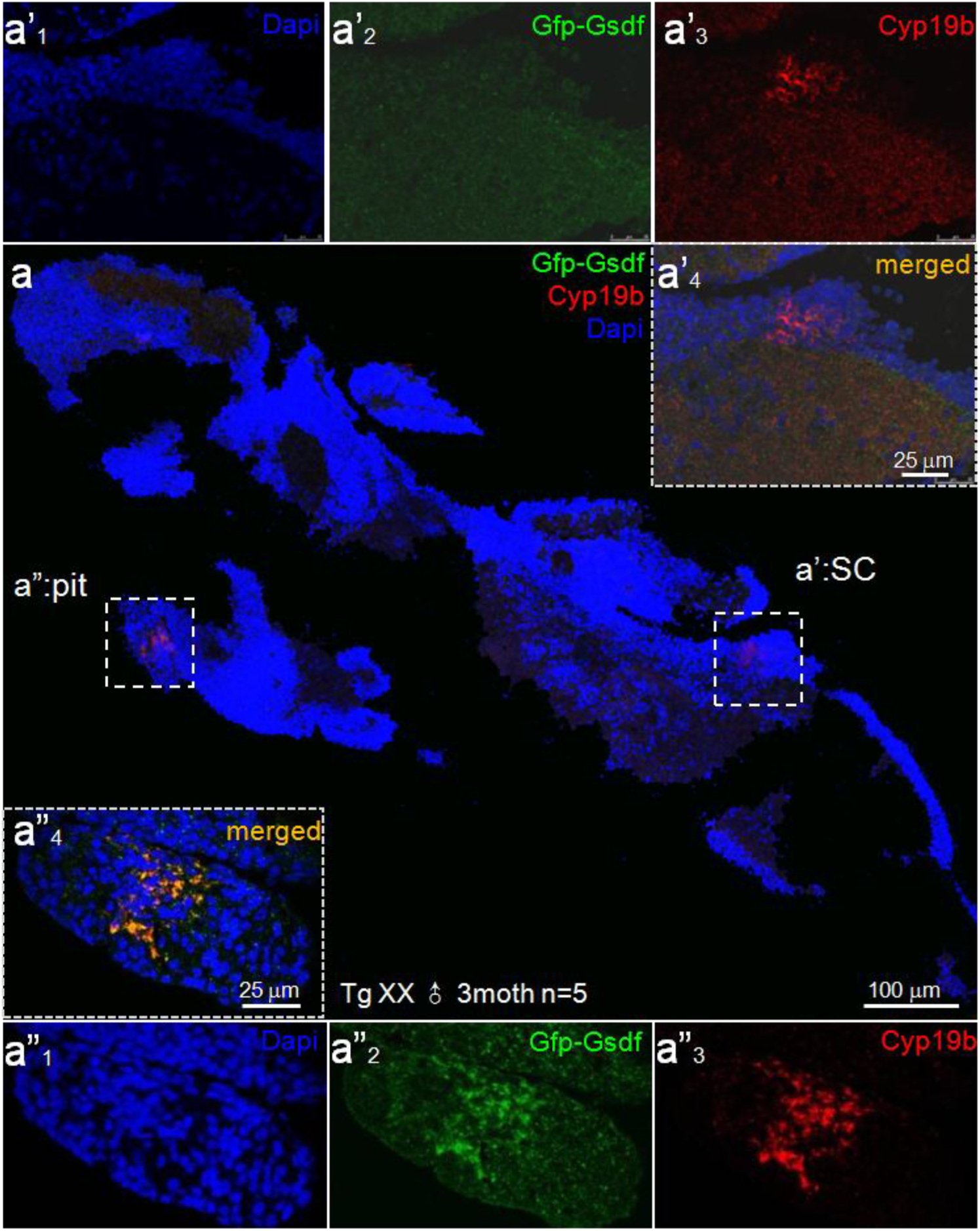
Transgenic *gsdf* expression co-localized with *cyp19b* in XX Tg male brain. (a) Immunocytohistological analysis revealed the expression of *cyp19b* and *gfp–gsdf* signals in the whole brain of Tg^cryG^ XX males. Cyp19b was detected in both pituitary (pit, a”) and spinal cord (SC, a’); Gfp was only defined in the pituitary (a” pit) in the longitudinal section of whole brain (n = 5 Tg brains examined). Magnified images of green Gfp and red Cyp19b expression, respectively, in the same sections. Nuclei with DAPI staining in blue.

**Figure S3.**
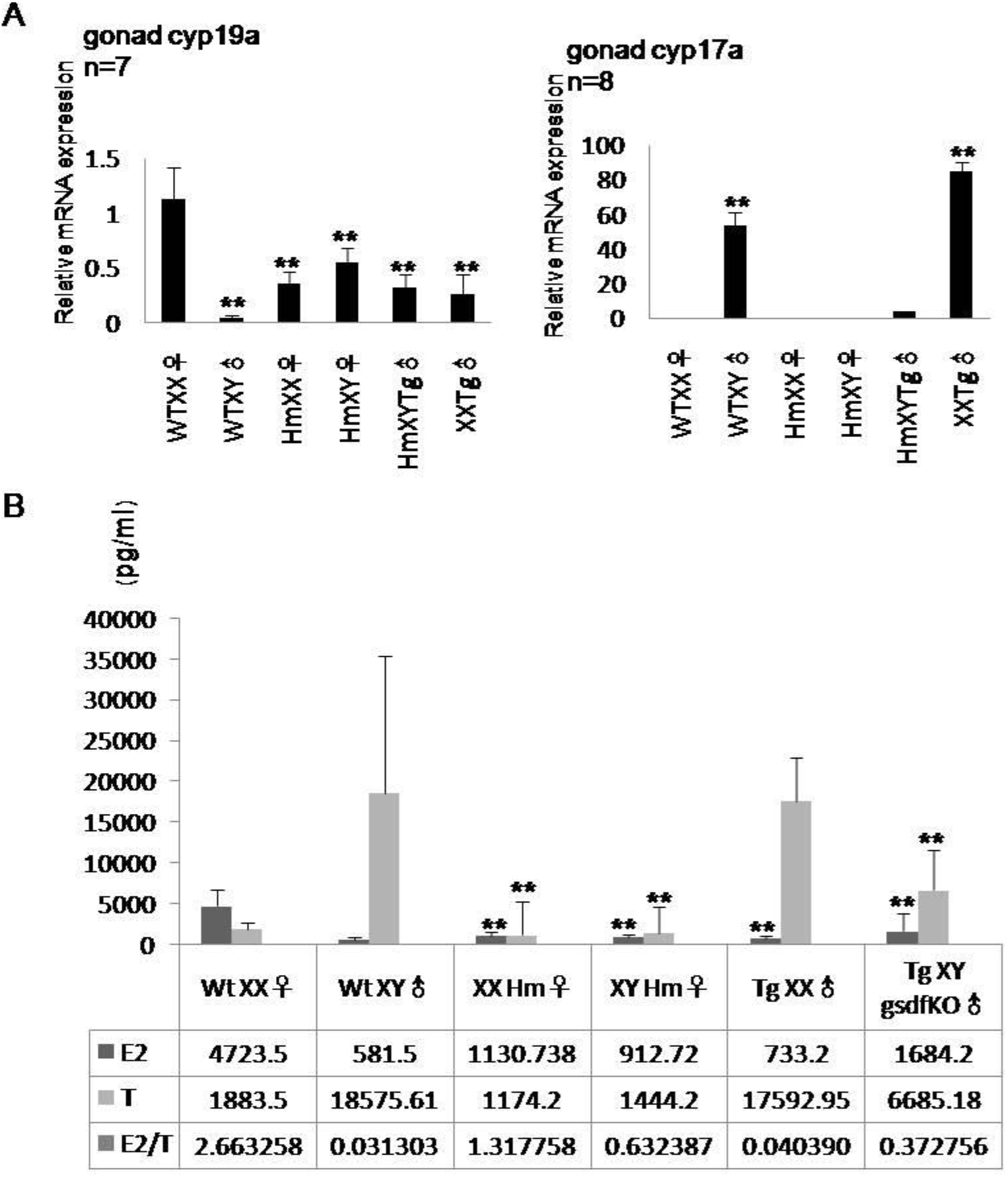
Expression profiles of relevant genes and E2/T levels in a variant line. (A) *cyp19a* and *cyp17a* are key enzymes in E2 and T production and reflect preferential expression in ovary or testis. (B) Ratio of serum E2/T lower in males and higher in females. All data represent means ± SEM from three measurements of individual serum mixtures (*n* = 6). ** *P* < 0.01.

